# Structure of the human RAD17-RFC clamp loader and 9-1-1 checkpoint clamp bound to a dsDNA-ssDNA junction

**DOI:** 10.1101/2022.03.11.484023

**Authors:** Matthew Day, Antony W. Oliver, Laurence H. Pearl

## Abstract

The RAD9-RAD1-HUS1 (9-1-1) clamp forms one half of the DNA damage checkpoint system that signals the presence of substantial regions of single-stranded DNA arising from replication fork collapse or resection of DNA double strand breaks. Loaded at the 5’-recessed end of a dsDNA-ssDNA junction by the RAD17-RFC clamp loader complex, the phosphorylated C-terminal tail of the RAD9 subunit of 9-1-1 engages with the mediator scaffold TOPBP1 which in turn activates the ATR kinase, localised through the interaction of its constitutive partner ATRIP with RPA-coated ssDNA.

Using cryogenic electron microscopy (cryoEM) we have determined the structure of a complex of the human RAD17-RFC clamp loader bound to human 9-1-1, engaged with a dsDNA-ssDNA junction. The structure answers the key questions of how RAD17 confers specificity for 9-1-1 over PCNA, and how the clamp loader specifically recognises the recessed 5’ DNA end and fixes the orientation of 9-1-1 on the ssDNA.

## INTRODUCTION

Substantial segments of single stranded DNA generated by stalled replication forks or resection of DNA double-strand breaks, trigger a DNA damage checkpoint response, which is dependent on activation of the PIKK-family protein kinase ATR and phosphorylation of its downstream target CHK1 (Guo et al., 2000; Sancar et al., 2004; Sperka et al., 2012) mediated by their mutual interaction with CLASPIN (Day et al., 2021b; Kumagai et al., 2004; Lindsey-Boltz et al., 2009). ATR is recruited to segments of ssDNA via interaction of its constitutive partner ATRIP (Cortez et al., 2001) with the replication protein A (RPA) complexes that coat ssDNA (Harrison and Haber, 2006; Itakura et al., 2004; Zou and Elledge, 2003).

Full catalytic activation of ATR at sites of ssDNA is dependent on the RAD9-RAD1-HUS1 (9-1-1) checkpoint clamp (Majka et al., 2006b) preferentially loaded at a 5’-recessed margin (Ellison and Stillman, 2003; Majka et al., 2006a) by a specialised form of the multi-protein Replication Factor C (RFC) clamp loader in which the large RFC1 subunit, required for loading the homotrimeric PCNA, is replaced by the smaller RAD17 (Bermudez et al., 2003; Majka and Burgers, 2004).

The presence of the 9-1-1 clamp loaded at a 5’-recessed dsDNA-ssDNA junction, is communicated to ATR localised on RPA-bound ssDNA, by the scaffold protein TOPBP1 (Day et al., 2021a; Garcia et al., 2005; Wardlaw et al., 2014), whose BRCT1 domain binds to a phosphorylated motif in the extended C-terminal tail of RAD9 (Day et al., 2018; Delacroix et al., 2007; Greer et al., 2003; Lee et al., 2007; Rappas et al., 2011; St Onge et al., 2003; Takeishi et al., 2010), and whose AAD-domain binds to and catalytically activates ATR (Kumagai et al., 2006; Mordes et al., 2008). In S and G2/M phases of the cell cycle this is sufficient for cell cycle checkpoint activation, however in the G1 phase an additional interaction with 53BP1 is required (Bigot et al., 2019).

We have now determined the cryoEM structure of the human RAD17-RFC2-RFC3-RFC4-RFC5 (RAD17-RFC) clamp loader complex bound to the 9-1-1 checkpoint clamp and engaged with a dsDNA-ssDNA junction. The structure explains how replacement of RFC1 with RAD17 changes the loading specificity of the RFC complex from PCNA to 9-1-1 and reveals the means by which the orientation of 9-1-1 loading at a dsDNA-ssDNA junction is dictated through recognition of the 5’ recessed end by RAD17.

## RESULTS

### Protein Expression and CryoEM

To obtain protein for structural analysis human 9-1-1 checkpoint clamp and RAD17-RFC clamp loader were expressed together in *Sf9* insect cells, and purified by affinity and size exclusion chromatography (see **METHODS**). The yields of assembled complex in initial trials were low due to the apparent lability of the human RAD17 protein, which was readily lost from the otherwise very stable RFC2-RFC3-RFC4-RFC5 clamp loader core. To circumvent this, we explored a fusion strategy in which RAD17 was genetically fused via an extended ‘linker’ to the N- or C-terminus of either RAD9, RAD1 or HUS1. While some fusions completely disrupted assembly, fusion of RAD17 to the C terminus of RAD1 significantly stabilised the overall complex, and delivered material in which all subunits co-purified, which co-migrated with a dsDNA-ssDNA junction in electrophoretic mobility assays, and that showed the presence of both clamp and clamp loader in negative stained electron micrographs (data not shown).

For cryoEM, the purified samples were mixed with ATP*γ*S and a DNA molecule that folds back on itself to give 24 base-pairs of fully complementary dsDNA capped with a tetraloop at one end, and with a 24 nucleotide 3’ overhang at the other (**see METHODS**). Samples were applied to grids and images collected on an FEI Titan Krios microscope equipped with a Falcon IV detector in counting mode (**see METHODS**). Movies were motion corrected and images processed in cryoSPARC (DiIorio and Kulczyk, 2022) and RELION 4.0 (Kimanius et al., 2021). The final map derived from 150626 particles, with an overall resolution of 3.59Å as defined by ‘gold standard’ Fourier shell correlation (Henderson et al., 2012), was post-processed using DeepEMhancer (Sanchez-Garcia et al., 2021). The image processing workflow is shown in **Figure S1**, and parameters of the data collection and refinement are given in **TABLE 1**.

**TABLE 1.**
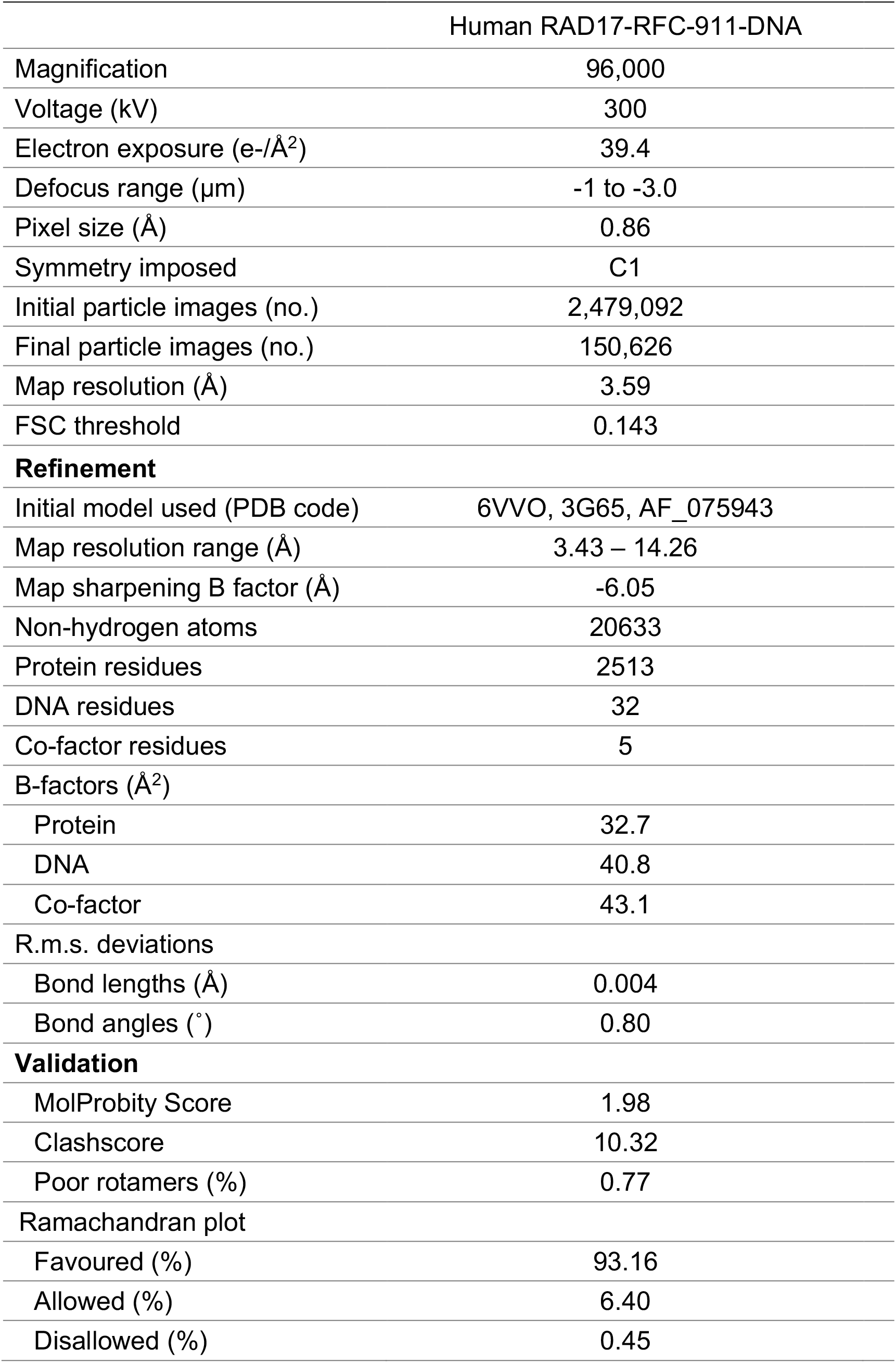

### Model Fitting

Preliminary inspection of the post-processed map confirmed the presence of both the 9-1-1 clamp and the RAD17-RFC clamp loader, as well a segment of dsDNA protruding from the core of the particle. As the visible duplex DNA and some clear *α*-helices had a left-handed thread, the map was inverted to the correct hand (**FIGURE 1A**).

**FIGURE 1.**
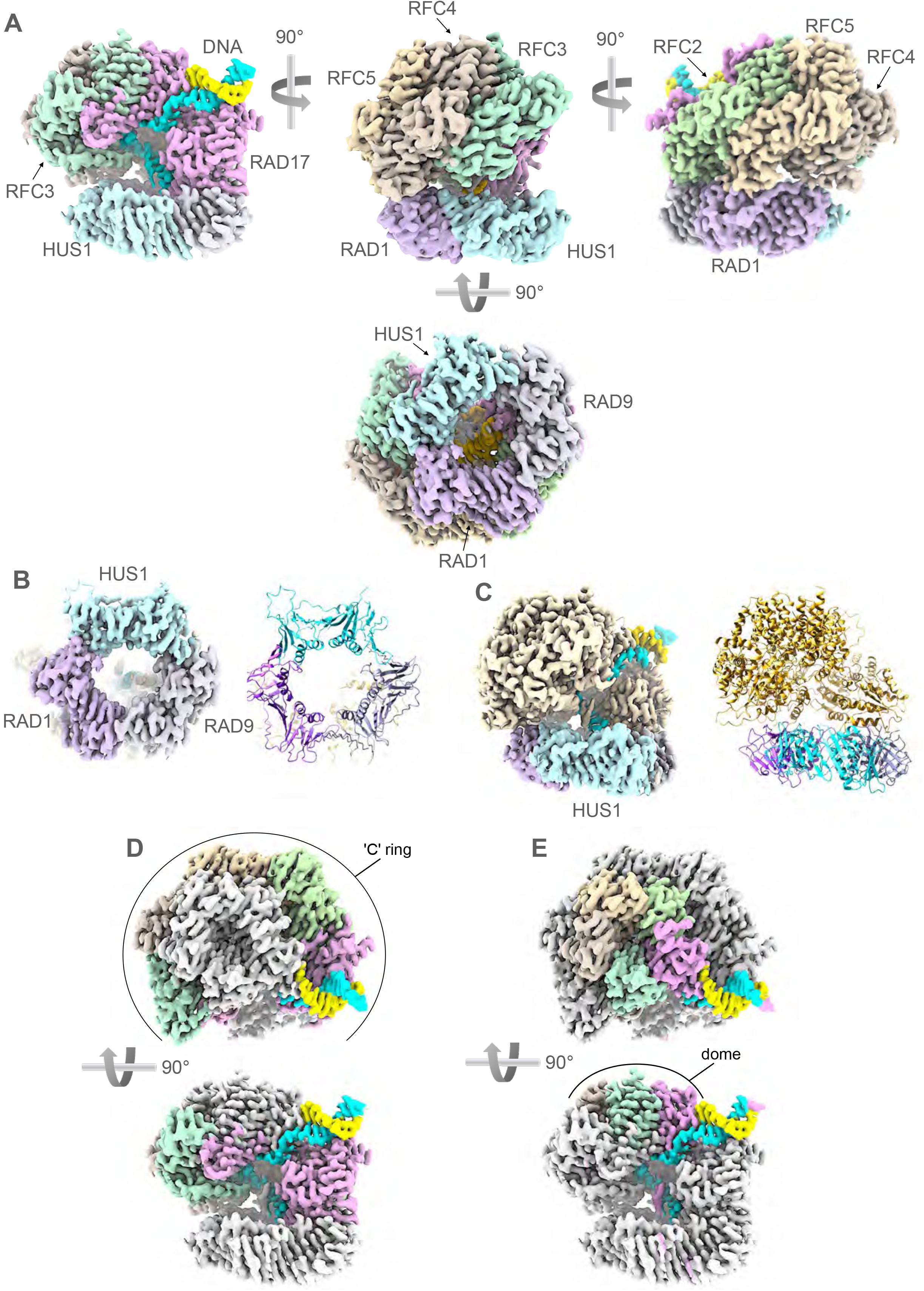
CryoEM structure of RAD17-RFC – 9-1-1 -DNA complex. **A**. Orthogonal views of the experimental map coloured to reflect the individual polypeptide chains. The two strands of the DNA are coloured sky blue and sunflower yellow – the colours of the Ukrainian flag. This and all other molecular graphics were created in ChimeraX (Pettersen et al., 2021) or Chimera (Pettersen *et al*., 2004). **B**. CryoEM volume for the 9-1-1 ring (left) and secondary structure of the fitted atomic model (right) showing that the ring is fully closed in this complex with RAD17-RFC. **C**. CryoEM volume for the complex ring (left) and secondary structure of the fitted atomic model (right) showing that the clamp-loader makes no contact with the HUS1 subunit of the 9-1-1 ring. **D**. Orthogonal views showing the Rossmann-fold and central helical bundle domains of RAD17, RFC2, RFC5, RFC4, and RFC (coloured as **A**) form a slightly twisted ‘C’ ring at the base of the clamp loader, that interacts with the 9-1-1 ring. **E**. As D. but highlighting the arrangement of the C-terminal domains of each subunit, that form a pentagonal dome at the top of the clamp loader.

An initial model for the 9-1-1 ring, which appeared fully closed, was obtained by placing the coordinates of our previously determined crystal structure of 9-1-1 (PDB code: 3G65) (Doré et al., 2009) into the map using Chimera (Pettersen et al., 2004). The correct orientation of the pseudo-threefold symmetric structure was determined by evaluating the fit to the map in three pseudo-equivalent orientations approximately 120°apart around the perpendicular to the ring, with both orientations of the faces of the ring. For RFC2, RFC3, RFC4 and RFC5, which each contain three globular domains connected by linker segments, initial models were obtained by manual docking of individual domains taken from a cryoEM structure of RFC1-RFC2-RFC3-RFC4-RFC5 in complex with PCNA (PDB code: 6VVO). For RAD17, for which no experimentally determined structure is available, a pared-down version of the predicted model (AF-075943-F1) in the EBI-AlphaFold database (Varadi et al., 2022) retaining only the high-confidence globular elements, was docked as two separate rigid bodies. Polypeptide segments linking the docked globular domains for all chains in the clamp loader were constructed using Coot (Emsley et al., 2010) into well-ordered features of the map. All chains are substantially complete with the exception of RAD17, where no place can be found within the experimental volume for several extended loops and terminal segments predicted to be disordered in the AlphaFold model of isolated human RAD17. Again, using Coot, an initial model for the bound DNA was obtained by docking a standard B-form DNA model into the map, and then manually constructing the ssDNA 3’ extension. The fit to the experimental volume of all the docked polypeptides chains was adjusted manually in Coot and the global fit optimised using phenix.refine (Afonine et al., 2018). Parameters defining the data collection and the quality of the final atomic model (PDB code: 7Z6H) are given in **TABLE 1**.

### Structure of the RAD17-RFC – 9-1-1 complex

The structure of 9-1-1 in the complex with RAD17-RFC is essentially identical to the crystal structure of the isolated clamp (Doré *et al*., 2009). All three subunits are coplanar, and all three interfaces between the subunits are closed and sealed by main-chain hydrogen bonding between the edge β-strands (**FIGURE 1B**). The ~120 residue extended C-terminus of RAD9 that carries the TOPBP1-binding motif, is not visible in the structure and is assumed to be completely disordered, as is the 45 residue linker connecting the C terminus of RAD1 and the N-terminus of RAD17.

The RAD17-RFC structure, sits over one face of the 9-1-1 ring, contacting the RAD9 and RAD1 subunits (see below) but making no contact with HUS1 (**FIGURE 1C**). RAD17-RFC consists of two effective layers. The lower part is an open and slightly twisted ‘C’ shaped ring formed by the N-terminal Rossmann fold domains and central helical bundle domains of RAD17, RFC2, RFC5, RFC4, and RFC3 (clockwise viewed from 9-1-1). A gap in the ring occurs between the Rossman and central domains of RAD17, and an additional C-terminal helical domain unique to RAD17 which packs against the Rossman domain of RFC3 (**FIGURE 1D**). The upper part of the structure is a closed pseudo-pentagonal ring formed by the C-terminal helical domains of the subunits in the same order (**FIGURE 1E**). The upper ring, which forms a ‘dome’ over the central cavity, is offset from the centre of the lower ‘C’ ring, lying predominantly over RFC3, RFC4 and RFC5.

Features corresponding to bound adenine nucleotides are identifiable in all five subunits of the clamp loader (**FIGURE S2**). While the conformations of the nucleotides differ, in all cases there is evidence for a *γ*-(S)-phosphate group being present, indicating that the state of the clamp-loader observed here corresponds to that prior to the hydrolysis of ATP.

### Clamp – clamp loader interactions

Most of the contacts between the clamp and the clamp loader, involve RAD1, which contacts RFC2, RFC5 and RFC4 (**FIGURE 2A**). The interaction with RFC2 is the least substantial of the three and occurs close to the edge of RAD1, involving an *α*-helix at residues 114-125 of RFC2 making weak polar and likely solvent mediated interactions in the shallow groove at the interface of RAD1 with RAD9 (**FIGURE 2B - left**). The adjacent clamp loader subunit RFC5, binds to a shallow depression near the middle of RAD1 adjacent to its C-terminal strand, with residues at the C-terminal end of helix 99-111 of RFC5 making predominantly polar interactions (**FIGURE 2B - middle**). RFC4 provides the third interaction with RAD1, close to its interface with HUS1 in the 9-1-1 ring, with RFC4 residues from the C-terminus of the *α*-helix at 116-126 and the following loop, making polar and likely solvent bridged interactions with the N-terminal end of the inter-domain linker (IDL) that connects the two halves of RAD1 (**FIGURE 2B - right**).

**FIGURE 2.**
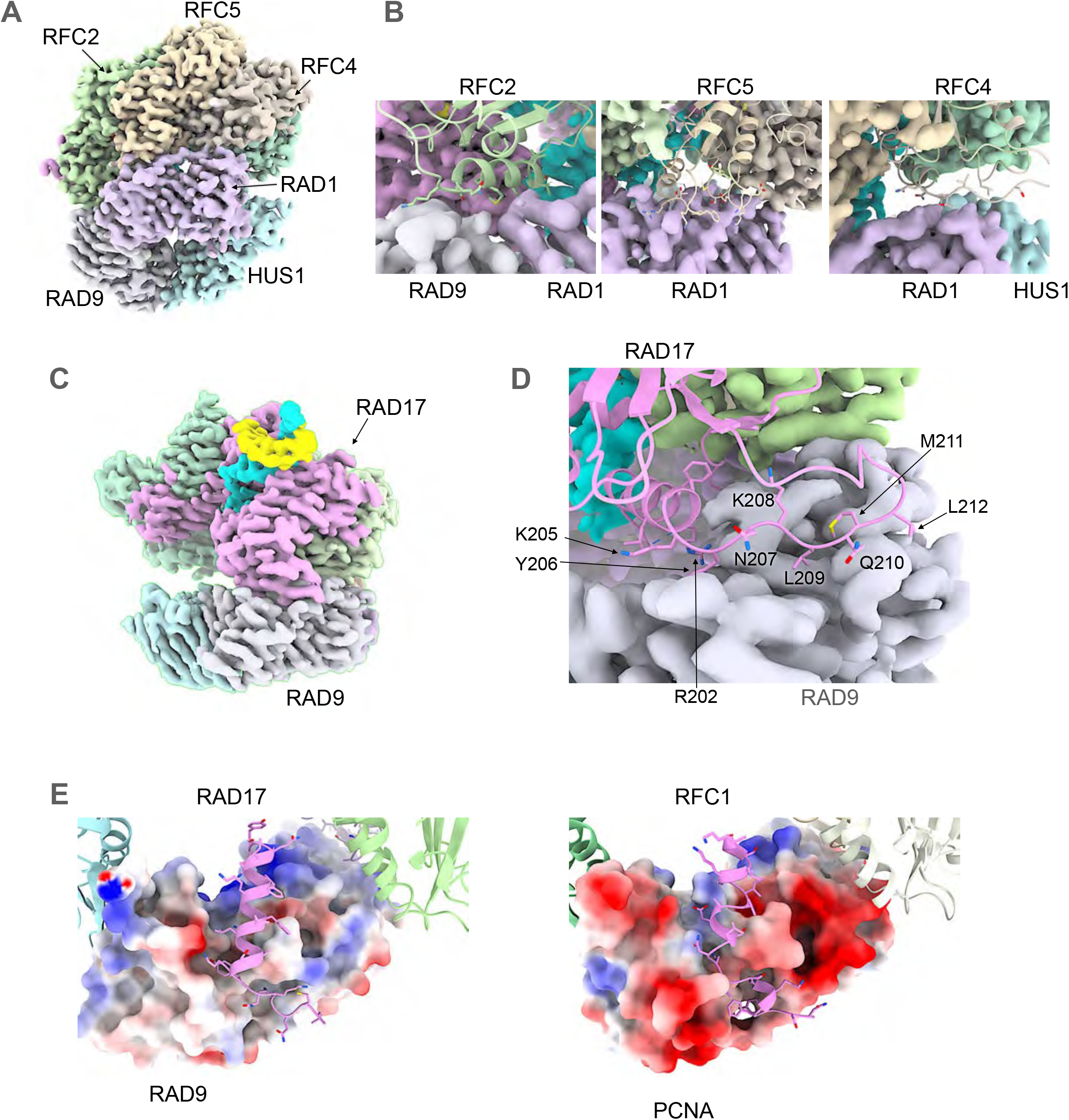
Clamp – clamp loader interactions. **A**. RAD1 interacts with RFC2, RFC5 and RFC4. **B**. Closeups of RAD1 interactions. The interfaces contain few intimate contacts and are likely mediated in part by solvent. The buried surface areas calculated by PISA (Krissinel and Henrick, 2007) are 114, 567 and 236Å^2^ respectively, suggesting that they do not play a major role in complex formation on their own. **C**. RAD17 interacts exclusively with RAD9. **D**. Detail of the RAD17 interaction with the surface of RAD9. The side chains of Leu209 and Met211, and of Tyr206 bind in hydrophobic pockets either side of the inter-domain linker of RAD9, while lysines 205 and 208, and Arg202, make polar interactions with RAD9 residues around the pocket rims. The buried surface area for this interface 1008Å^2^ suggests it makes a substantial contribution to the overall interaction between the clamp and the clamp loader. **E**. Comparison of the interactions between RAD17 and RAD9 (left) with those between RFC1 and PCNA (right). Although the overall topology of the interacting regions are similar, multiple detailed differences in the interfaces make the two clamp loader components mutually incompatible with the other’s preferred clamp component and ensure their exclusivity and specificity.

The other major interface between the clamp and clamp loader is provided by the RAD17 and RAD9 subunits **(FIGURE 2C)**. This is the most substantial contact between the two complexes and is mediated by the interaction of a linear segment of RAD17 – an *α*-helix 194-205 and the following strand to 212 – which bind into two distinct connected pockets close to the junction of the two halves of RAD9. The core of the interaction is provided by Tyr206 of RAD17 plugging into a pocket in RAD9 lined by the side chains of Ser46, Glu165, Cys218, Ala263, and Leu265, and the peptide backbone of Phe217 and Thr264, and by Leu209 and Met211 of RAD17, which together fill a pocket in RAD9 lined by Val41, Arg45, Ala47, Tyr48, Leu132, Gln133, Ala134, Val135, Val261, and Ala263, positioned in the equivalent position to the PIP box binding pocket in PCNA. Both of these pockets had previously been noted as likely binding sites for interacting proteins in the original crystal structure of 9-1-1 (Doré *et al*., 2009) (**FIGURE 2D**). These core hydrophobic interactions are supported by a plethora of polar interactions, and a high degree of surface charge complementarity, especially between the strongly positive region of RAD17 surrounding Tyr206 (including Lys205), and the correspondingly strongly negative regions surrounding the pocket in RAD9 that accommodates it. Consistent with our observations, deletion or mutation of this highly conserved 205-KYxxL-209 motif in RAD17 has been shown to abolish interaction of RAD17 and RAD9 *in vivo* (Fukumoto et al., 2016).

### Structural basis of clamp selection

The RFC2, RFC3, RFC4 and RFC5 subunits of the RAD17-RFC clamp loader complex, are also functionally associated in the canonical RFC clamp loader for the replicative processivity factor PCNA (Kelch et al., 2012) where RAD17 is substituted by the structurally related RFC1. Comparison of the surfaces of homotrimeric human PCNA with the heterotrimeric human 9-1-1 complex (Doré *et al*., 2009) revealed substantial differences that would require complementary and distinct interactions by the two different assemblies that mediate their loading. In order to understand these, we have compared the structure presented here with the cryoEM structure of a human RFC-PCNA complex (PDB code 6VVO), (Gaubitz et al., 2020).

Both RAD17 and RFC1 make the most substantial interaction in their respective complexes, involving a linear motif 189-212 in RAD17 and 683-705 in RFC1 (**FIGURE 2E,F**) interacting with the surface of RAD9 or one of the three identical PCNA molecules, respectively. In both cases, the interaction involves insertion of several hydrophobic residues from the clamp loader component into pockets either side of the IDL. However, the geometry of the interaction in each case is specific and exclusive, ensuring specificity for the particular species of clamp the loader engages with. Thus, PCNA lacks the required pocket to accommodate Lys205 and Tyr206 in RAD17, with the corresponding pocket in PCNA, lined with Lys254, Val45, Ala252, Pro253, Ala208, Tyr211, being shallower than the one in RAD9 and perfectly ordered to instead accommodate two asparagine residues, Asn695 and Asn696, in the corresponding sequence in RFC1. The second half of the clamp loader linear motif interaction for RFC1 and PCNA uses a canonical PIP binding motif in RFC1 that is lacking in the alternative RAD17 subunit, likewise the corresponding pocket to the PIP box binding pocket in RAD17 is shorter than required for canonical PIP binding.

Additional to the interactions of the clamp-specific components which likely drive complex formation, the common components of RFC2 and RFC5 make less substantial interactions with components of the clamp in both clamp loader settings. Thus, RFC2 uses essentially the same side chains in the helical segment from 114-127 to contact both RAD1 and PCNA in the different complexes but does so in subtlety different ways. Similarly, residues 98-115 of RFC5 adapt to interactions with RAD1 or PCNA by utilising different residues within the segment. In both cases the interacting surfaces are much smaller than the substantial interfaces made with RAD17 or RFC1 and would play no significant role in mediating specificity.

### DNA binding and ds-ssDNA junction recognition

Contrary to models based on the seminal crystal structure of the RFC-PCNA clamp loader – clamp complex (Bowman et al., 2004) in which DNA was proposed to run through the centre of the ring formed by the RFC subunits and interact with them all, the DNA in the complex presented here, enters the RAD17-RFC – 9-1-1 complex from the side, interacting solely with the RAD17 subunit (**FIGURE 3A**). The bound segment of dsDNA interacts with the N-terminal Rossman fold and following 3-helix bundle domains of RAD17, with polar and charged residues 288-292 from the edge of the central β-sheet, and at the N-terminal ends of helices 292-307 and 106-121 interacting with the sugar-phosphate backbone and minor groove of the DNA (**FIGURE 3B**). The conformation of the DNA in this region is predominantly canonical B-form, but with some slippage and twist of the base pairs evident.

**FIGURE 3.**
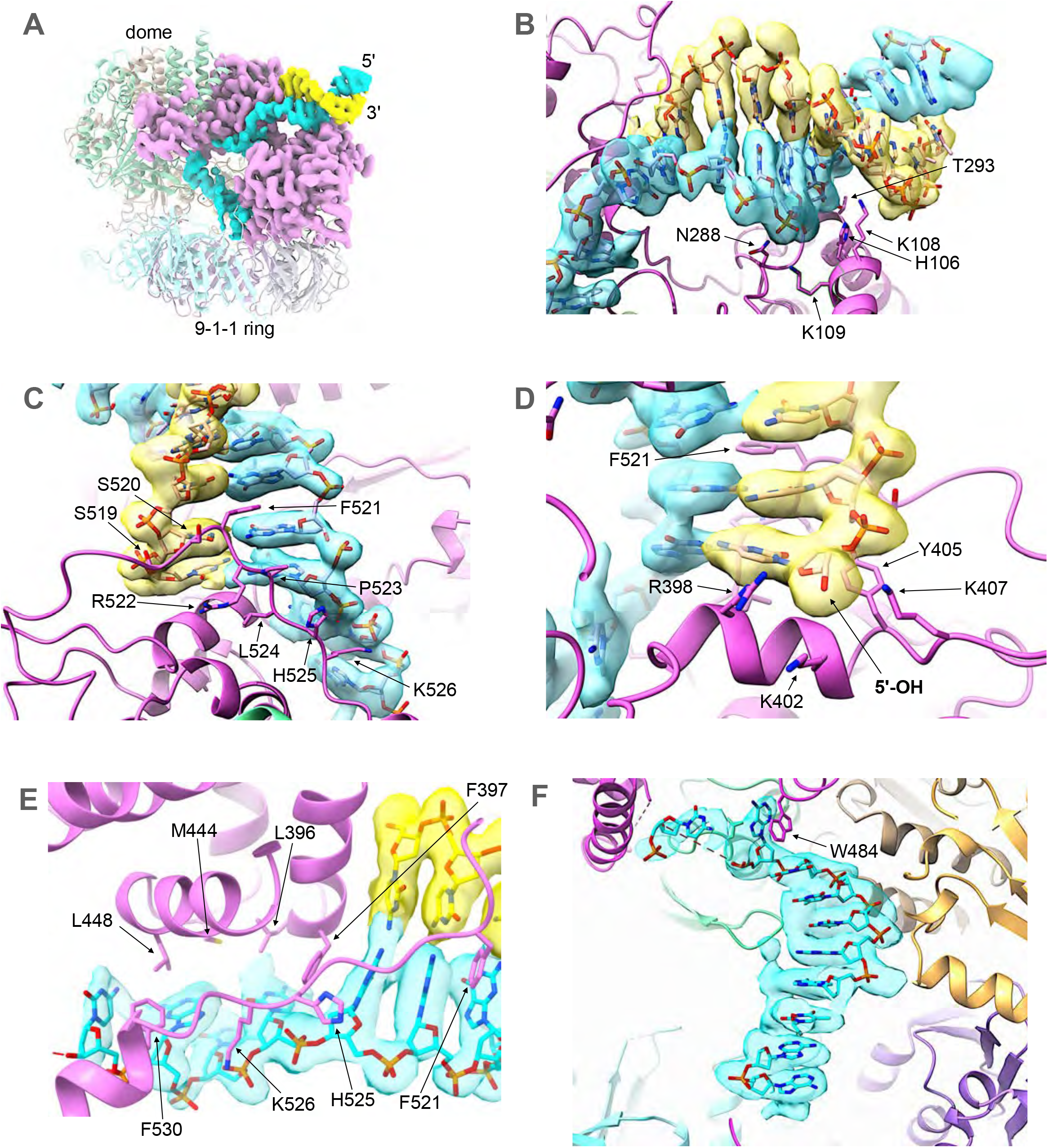
DNA binding by the clamp loader. **A**. The double stranded DNA (blue and yellow) enters the clamp loader from the side, interacting exclusively with RAD17, and pushing through a gap between the N-terminal Rossman and central domains of RAD17, and its C-terminal helical domains. The path of the single stranded segment of DNA is twisted by ~90°as it passes under the dome, to direct it towards the 9-1-1 ring bound at the base of the clamp loader. **B**. Details of the interaction of the N-terminal and central domains of RAD17 with the dsDNA. Multiple basic and polar residues interact with the sugarphosphate backbone of the duplex DNA, in and around the minor groove. The experimental volume for the DNA is shown as a transparent surface through which the atomic model can be seen. **C**. Details of the interaction of the C-terminal domains of RAD17 with the dsDNA. As well as polar interactions with the sugar phosphate backbones of both strands, the side chain of Phe521 partially intercalates into the DNA through the minor groove, disrupting base pairing and stacking. **D**. The path of the 3’-5’ strand is blocked by an *α*-helix from RAD17, with the side chains of three basic residues Arg398, Lys402 and Lys407, forming a pocket into which the 5’-OH of the interrupted strand, projects. This pocket would be able to bind and neutralise a 5’-phosphate if one were present. **E**. The path of the 5’-3’ strand is twisted away from the dsDNA-ssDNA junction over four nucleotides through interaction of their sugar-phosphate backbones with multiple charged residues, and the exposed faces of their bases with an arc of hydrophobic side chains. **F**. A further twist in the path of the ssDNA is dictated by interaction with the side chain of Trp484, before a canonical B-DNA like conformation is re-established as the remaining visible ssDNA spirals down into the central hole of the 9-1-1 ring.

The continuity of the duplex DNA as it runs into the body of the complex, is disrupted by a segment of RAD17 from 519-527, which runs along the outside of the minor groove of the DNA, interacting with the sugar-phosphate backbone of both strands. The side chain of Phe521 in this segment partially intercalates into the base-pair stack, disrupting the complementary hydrogen bonding either side of it (**FIGURE 3C)**. Complete disruption of the duplex structure occurs two base pairs further on, where the 5’ end of one of the strands runs into the side of an *α*-helix 395-405 that blocks any further passage (**FIGURE 3D**). The deoxyribose sugar and associated base of the 5’ nucleotide on the interrupted strand sits on a platform formed by the peptide of Gly401 and the side chain of Tyr405, with its terminal 5’-hydroxyl directed into a narrow pocket defined by the basic side chains of Arg398, Lys402 and Lys407. In the DNA used here, the 5’-end is not phosphorylated, but a 5’-phosphate could be readily accommodated by this pocket, where the positive charge of the basic side chains would neutralise the charge on the terminal phosphate.

The uninterrupted DNA strand running in a 5’ to 3’ direction away from the ss-dsDNA junction, is twisted towards the centre of the cavity under the ‘dome’ through interactions with the 519-530 segment, and residues from *α*-helices 446-456 and 396-405. The first four nucleotides of this ssDNA segment which can be readily placed into the cryoEM volume, adopt a very non-canonical conformation, driven by association of the unstacked faces of their bases, with a hydrophobic arc formed by the side chains of Phe530, Leu448, Met444, Leu396 and Phe397 (**FIGURE 3E)**. After this the map fades out, suggesting that the ssDNA becomes conformationally disordered at this point. However, further into the cavity, after a gap of two or three nucleotides, the ssDNA becomes ordered again, with the first two nucleotides of this segment stacking parallel and perpendicularly respectively with the indole side chain of Trp484 of RAD17, which sits at the tip of a loop connecting helices 464-481 and 486-505. A further seven nucleotides are clearly visible running as a single helix almost perpendicular to the direction of the duplex DNA, down towards the hole at the centre of the 9-1-1 ring (**FIGURE 3F**). A weak isolated feature suggestive of polypeptide is visible interacting with the edges of the bases in the ssDNA segment. This is likely to be part of a predicted disordered loop in RAD17 from 167-189 which would project into the main cavity of the complex, but the detail is insufficient to map it to the RAD17 sequence.

## DISCUSSION

The structure presented here explains how the incorporation of RAD17 in place of RFC1 switches the specificity of the clamp loader from the homotrimeric PCNA with its myriad roles in DNA replication and repair, to the heterotrimeric RAD9-RAD1-HUS1 clamp whose only well-established role is as one half of the DNA damage checkpoint system that signals under-replicated DNA. In that context, the structure also shows that RAD17 provides essentially all the interactions with the DNA, and most importantly, confers the specificity for a recessed 5’-end that arises when DNA replication has been interrupted, to enable 9-1-1 to form one half of the ‘two-factor’ trigger mechanism for the ATR/CHK1 arm of the DNA damage response (Bartek and Lukas, 2003).

With the checkpoint clamp localised at the recessed 5’-end by its association with the clamp loader, the extended C-terminal tail of RAD9 is then available to bind to TOPBP1, itself localised to the adjacent ssDNA through its interaction with ATR-ATRIP bound to RPA (**FIGURE 4A**). The architecture of this larger complex and whether the two halves of the trigger are just loosely tethered by mutual interaction with TOPBP1 or form a compact super-complex, is currently unknown. Favouring the latter, loading of RAD17-RFC - 9-1-1 at dsDNA-ssDNA junctions appears to be stimulated by RPA (Zou et al., 2003), and a direct interaction between a motif in the C-terminal tail of RAD9 with a subunit of RPA has also been described (Xu et al., 2008).

**FIGURE 4.**
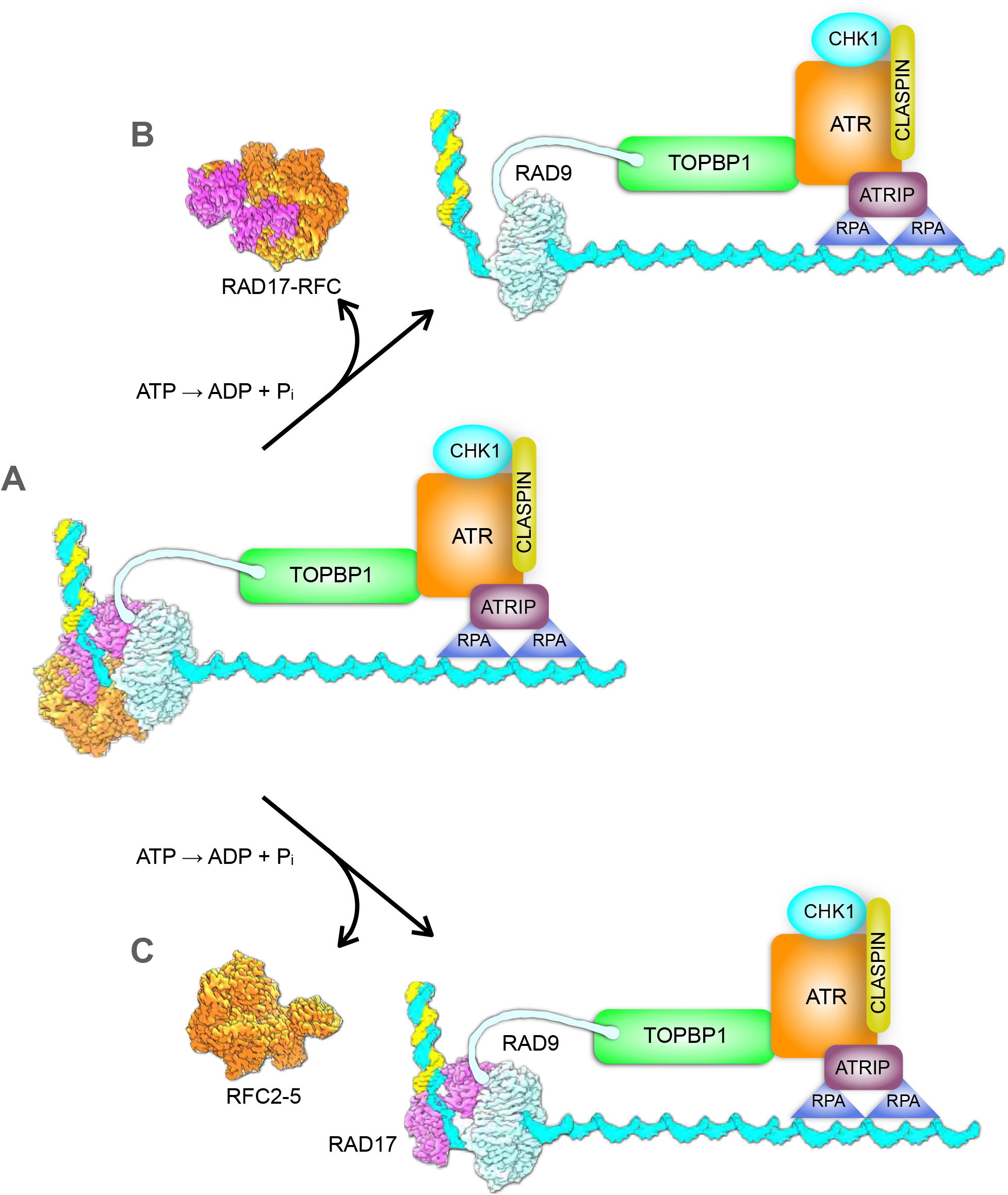
DNA damage checkpoint signalling. **A**. The structure described here provides a model for how the RAD17-RFC clamp loader orientates the 9-1-1 ring at a dsDNA-ssDNA junction to face the ssDNA and thereby interact with TOPBP1 and the other part of the DNA damage checkpoint signalling super-complex, prior to hydrolysis of the ATP bound to the components of the clamp loader. However, it is not clear what role subsequent ATP hydrolysis might play. **B**. ATP hydrolysis could promote release of the clamp loader from the clamp, leaving the 9-1-1 potentially free to diffuse away from the other components of the checkpoint super-complex, unless the critical interaction with TOPBP1 has been established. **C**. Alternatively, ATP hydrolysis could promote release of only the constitutively associated RFC2-5 sub-complex, which would be free to acquire a replacement RAD17 or more likely RFC1, which is present at substantially higher abundance (Wang et al., 2015). In this model, RAD17 would help retain 9-1-1 at the dsDNA-ssDNA junction and in proximity to the other components of the checkpoint super-complex to promote ATR and CHK1 activation.

Although all five of the subunits of RAD17-RFC, as with RFC1-RFC, can bind and are in principle capable of hydrolysing ATP, the role of this in the clamp loading process is still poorly understood (Gaubitz *et al*., 2020; Kelch *et al*., 2012). It is generally agreed that ATP (or ATPγS) binding is required for association of 9-1-1 with the clamp loader (Bermudez *et al*., 2003) and consequently favours its ultimate deposition on DNA (Zou *et al*., 2003). What role hydrolysis of that ATP plays is far from clear, and the apparent ATPase activity of RAD17-RFC is in any event, low compared to the canonical RFC1-containing complex that loads PCNA (Ellison and Stillman, 2003). In the structure presented here, in which all five binding sites are occupied by ATPγS and therefore representative of a pre-hydrolysis state, the clamp loader is properly positioned on the dsDNA-ssDNA junction and the clamp itself is fully closed encircling ssDNA and orientated so that the tail of RAD9 can engage with the other components of the checkpoint super-complex anchored to the ssDNA. What then would subsequent ATP hydrolysis achieve?

One possibility is that ATP hydrolysis facilitates the release of the clamp from the clamp loader, as has been shown for release of PCNA from the RFC1-RFC complex (Chen et al., 2009). However, unlike PCNA which is loaded in multiple copies and freely migrates on dsDNA as the anchor for a plethora of ‘client’ proteins involved in DNA replication and repair (Moldovan et al., 2007), only a single copy of 9-1-1 loaded on to ssDNA may be required (Bermudez *et al*., 2003), and it is far from clear that the RAD17-RFC clamp loader needs to fully detach from the loaded clamp for it to perform its checkpoint role.

Full dissociation of RAD17-RFC from the loaded 9-1-1 could even be deleterious, as it would allow the clamp to diffuse away from the loading site, losing its ‘context’ and the proximity to TOPBP1, ATR-ATRIP and RPA-coated ssDNA that is required to fire the DNA damage checkpoint (**FIGURE 4B**). As RAD17 provides the entirety of the interaction surface with the bound DNA, as well as making a substantial interface with RAD9 that would not obviously be disrupted by changes in conformation driven by ATP-hydrolysis – either in RAD17 or the generic RFC components – an alternative model is possible. In this, ATP hydrolysis would promote release of the stable, constitutively assembled, and functionally generic RFC2-RCF3-RFC4-RFC5 subcomplex, leaving the more loosely associated exchangeable RAD17 subunit still bound to 9-1-1 and maintaining its localisation at the dsDNA-ssDNA junction (**FIGURE 4C**). This model could account for the observation of RAD17 being phosphorylated by activated ATR (Bao et al., 2001) and for reports of its direct interaction with other DNA-bound factors such as the NBS1 component of the MRN complex (Wang et al., 2014). Further work will be required to determine which of these models is correct.

## NOTE

While this work was in progress, pre-prints describing structural studies of the homologous Rad24-RFC - Rad17-Mec3-Ddc1 complex from the yeast *Saccharomyces cerevisiae* were deposited in bioRxiv (10.1101/2021.10.01.462756 and 10.1101/2021.09.13.460164). As atomic coordinates for these structures are not publicly available at the time of writing, we have not been able to undertake a detailed comparison of the human and yeast complexes. However, the key features we describe here appear to be substantially conserved in the yeast system.

## ACKNOWLEDGMENTS

We thank Rebecca Thompson and Daniel Maskell for assistance with cryoEM data collection, Pascale Schelleberger and Fabienne Beuron for assistance and advice regarding grid preparation and evaluation, and Mathieu Rappas and Charles Grummitt for contributions at earlier stages of this work. This work was supported by Cancer Research UK Programme Grants C302/A14532 and C302/A24386 (A.W.O and L.H.P.).

## AUTHOR CONTRIBUTIONS

Conceptualisation: M.D., A.W.O., L.H.P.; Methodology: M.D., A.W.O., L.H.P.; Validation: M.D., A.W.O., L.H.P.; Formal Analysis: M.D., A.W.O., L.H.P.; Investigation: All Authors; Writing – Original Draft: L.H.P.; Writing – Review & Editing:M.D., A.W.O., L.H.P.; Visualisation: L.H.P.; Supervision: L.H.P., A.W.O.; Funding Acquisition: L.H.P., A.W.O.

## METHODS

### Protein Expression and Purification

RAD17RFC-9-1-1, was produced by infecting *Sf9* cells with a baculovirus coding for a construct containing RAD9-HIS, RAD1-STREP-RAD17-HIS HUS1 RFC2 RFC3 RFC4 HIS-RFC5 produced using the biGBac system (Weissmann et al., 2016). Cell pellets were re-suspended in lysis buffer containing 25mM HEPES pH 7.5, 200mM NaCl, 0.5mM TCEP, and supplemented with 10 U DNASE Turbo (Thermo Fisher Scientific) and cOmplete, EDTA-free Protease Inhibitor Cocktail (Merck), then disrupted by sonication, and the resulting lysate clarified by centrifugation at 40,000 x g for 60 minutes at 4°C. The supernatant was applied to a 5ml HiTrap TALON crude column (GE Healthcare) washed first with lysis buffer, followed lysis buffer supplemented with 10 mM imidazole, with retained protein then eluted by application of the same buffer but now supplemented with 250mM imidazole. The eluted protein was diluted with lysis buffer before application to a 5ml HiTrap STREP column (GE Healthcare), washed with lysis buffer and eluted using buffer supplemented with 2 mM Desthiobiotin. Following concentration, the sample was mixed with a 100 x excess of ATP*γ*S (Sigma) and applied to a Superdex200 10/300 size exclusion column (GE Healthcare) to purify the protein to homogeneity in 10mM HEPES pH 7.5, 150 mM NaCl, 0.5mM TCEP. DNA at a concentration of 1 mM with the sequence CCCGTATATTCTCCTACAGCACTAAATAATAGTGCTGTAGGAGAATATACGGGCTGCTCGTGTTGACAAGTACTGAT (IDT) was annealed to produce a hairpin DNA molecule with 24 base-pairs of fully-complementary dsDNA capped with a tetraloop at one end, and with a 24 nucleotide 3’ overhang. Peak fractions were mixed with a further 100 x excess of ATP*γ*S (Sigma) and a 1.2 x excess of the hairpin DNA and left at 4°C for 16 hours.

### Electron Microscopy and Image Processing

Quantifoil 1.2/1.3, 300 mesh copper grids (Quantifoil) were glow discharged using a Tergeo Plasma Cleaner (Pie Scientific) with an indirect plasma treatment for 30 seconds. Grids were loaded into a Leica EM GP2 (Leica microsystems) and 3 μl of a 600 nM sample was applied to the front of the grid, with an additional 0.5 μl buffer applied to the grids rear, before back blotting for 4 seconds and plunging into an ethane propane mix. Grids were stored in liquid nitrogen prior to imaging at 300 kV on a FEI Titan Krios (Thermo Fisher Scientific) equipped with Falcon 4 detector (Thermo Fisher Scientific). 8271 movies were collected, using data acquisition software EPU (Thermo Fisher Scientific), at a magnification of 96000 and a pixel size of 0.86Å using a total dose of 39.4 e-/Å2. These were motion corrected in 5×5 patches and the Contrast Transfer Functions (CTF) of the resulting micrographs calculated using MotionCor2 (Zheng et al., 2017) and CTFFIND4 (Rohou and Grigorieff, 2015) implemented in RELION4.0 (Kimanius *et al*., 2021). All subsequent processing used RELION4.0. Automated picking using the Laplacian of Gaussian method in RELION4.0 yielded 2479092 particles, which were extracted with 3x-binning and subjected to 2D classification. Of the initial picks, 304686 showed projections with clear features of the target complex and were subjected to further cycles of 2D classification to eliminate poor quality particles. An initial model calculated with a 10% subset of the 235039 retained particles, gave a promising volume, which was then used for several cycles of 3D classification with low resolution and/or poorly defined classes eliminated to yield a set of 155676 particles.

These were 3D refined and sharpened to give a model with a resolution of 5.16Å estimated by ‘gold standard’ Fourier Shell Correlation (Henderson *et al*., 2012). Refined particles were re-extracted at their full pixel size and subjected to single rounds of 2D and 3D classification resulting in a final set of 150626 particles. 3D refinement and post-processing of these gave a volume at 4.71Å resolution. CTF refinement and particle ‘polishing’ improved the map to 3.80Å, and further cycles of CTF and orientational refinement improved this further to 3.59Å. Subsequent 3D reclassification of these particles without orientational refinement failed to identify any distinct sub-classes with superior resolution or visual appearance, suggesting that the dataset was homogeneous. Finally, the un-sharpened map calculated from the refined particles was subjected to a deep learning-based sharpening and density modification process using DeepEMhancer (Sanchez-Garcia *et al*., 2021) to give the map used for model building and interpretation. A flow-diagram of the processing workflow is given in **FIGURE S1** and parameters of the data collection and refinement are given in **TABLE 1**.

### Data Deposition

CryoEM maps and refined coordinates have been deposited in the EMDB and Protein Databank as EMD-14527 and PDB ID 7Z6H respectively.

---

**SUPPLEMENTARY FIGURE 1.**
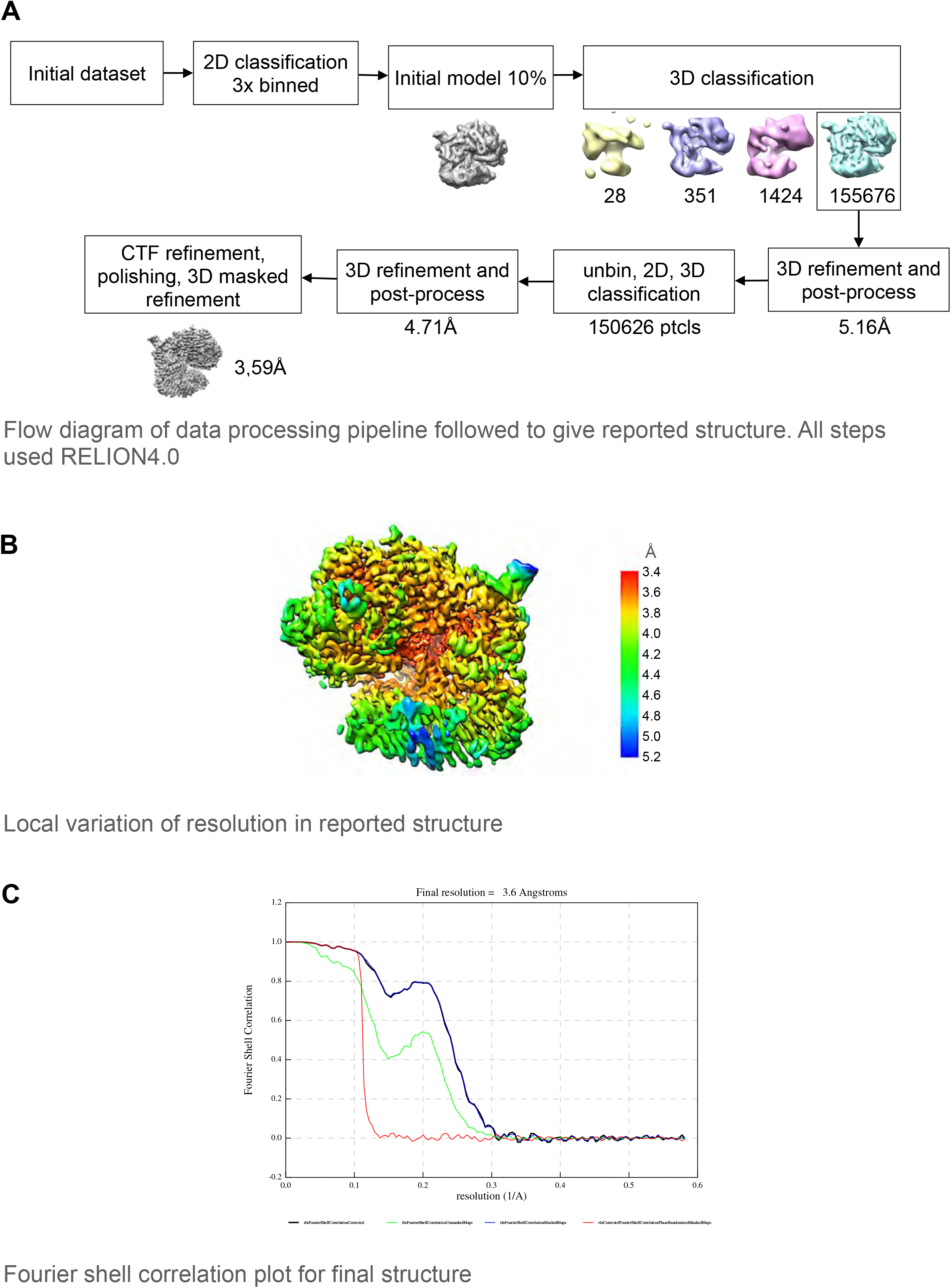

**SUPPLEMENTARY FIGURE 2.**
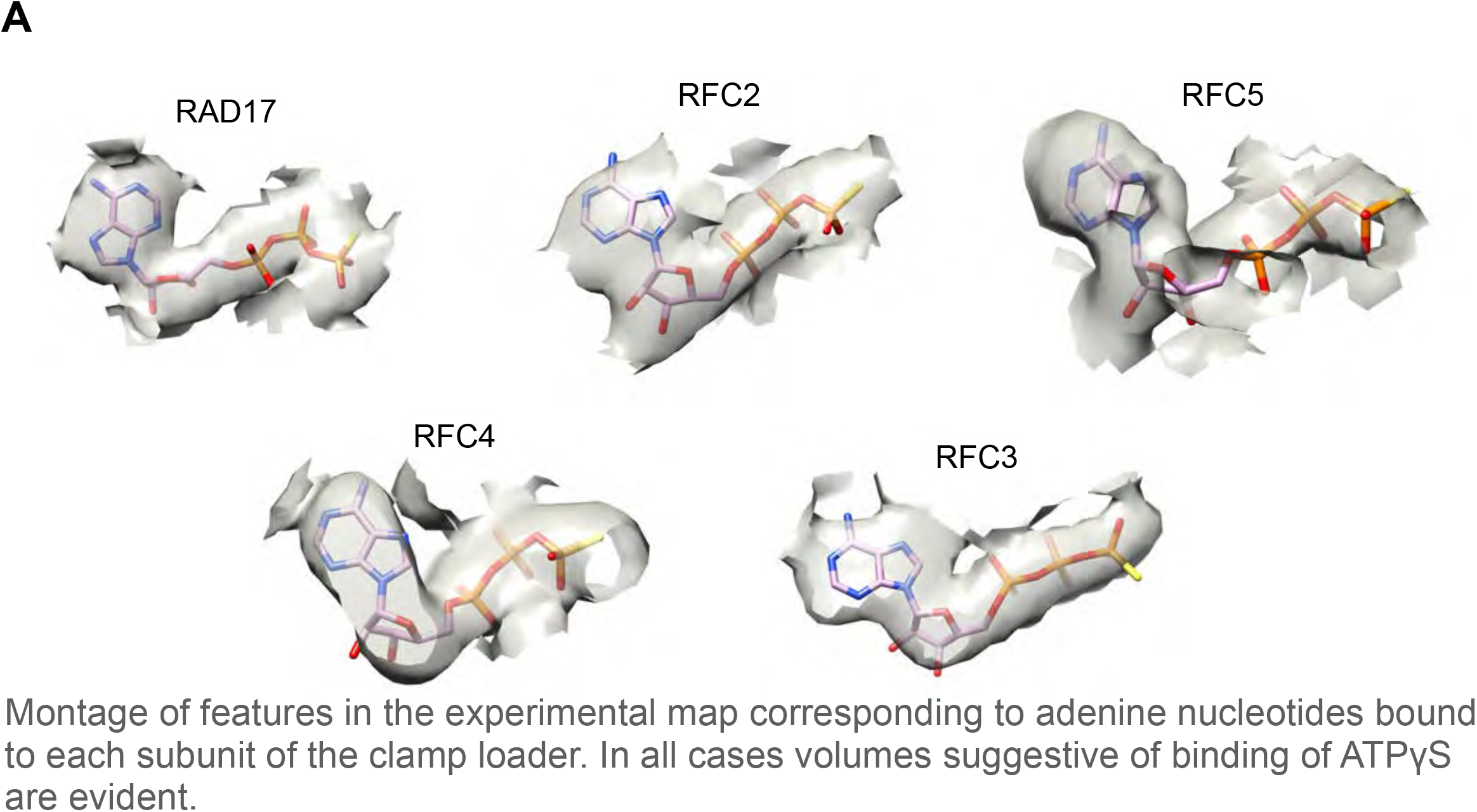

